# 11β-HSD1 inhibition does not affect murine tumour angiogenesis but may exert a selective effect on tumour growth by modulating inflammation and fibrosis

**DOI:** 10.1101/2021.08.04.455121

**Authors:** Callam T Davidson, Eileen Miller, Morwenna Muir, John C. Dawson, Martin Lee, Stuart Aitken, Alan Serrels, Scott P Webster, Natalie Z. M. Homer, Ruth Andrew, Valerie G. Brunton, Patrick W. F. Hadoke, Brian R. Walker

## Abstract

Glucocorticoids inhibit angiogenesis by activating the glucocorticoid receptor. Inhibition of the glucocorticoid-activating enzyme 11β-hydroxysteroid dehydrogenase type 1 (11β-HSD1) reduces tissue-specific glucocorticoid action and promotes angiogenesis in murine models of myocardial infarction. Angiogenesis is important in the growth of some solid tumours. This study used murine models of squamous cell carcinoma (SCC) and pancreatic ductal adenocarcinoma (PDAC) to test the hypothesis that 11β-HSD1 inhibition promotes angiogenesis and subsequent tumour growth. SCC or PDAC cells were injected into female FVB/N or C57BL6/J mice fed either standard diet, or diet containing the 11β-HSD1 inhibitor UE2316. SCC tumours grew more rapidly in UE2316-treated mice, reaching a larger (P<0.01) final volume (0.158 ± 0.037 cm^3^) than in control mice (0.051 ± 0.007 cm^3^). However, PDAC tumour growth was unaffected. Immunofluorescent analysis of SCC tumours did not show differences in vessel density (CD31/alpha-smooth muscle actin) or cell proliferation (Ki67) after 11β-HSD1 inhibition, and immunohistochemistry of SCC tumours did not show changes in inflammatory cell (CD3- or F4/80-positive) infiltration. In culture, the growth/viability (assessed by live cell imaging) of SCC cells was not affected by UE2316 or corticosterone. Second Harmonic Generation microscopy showed that UE2316 reduced Type I collagen (P<0.001), whilst RNA-sequencing revealed that multiple factors involved in the innate immune/inflammatory response were reduced in UE2316-treated SCC tumours.

11β-HSD1 inhibition increases SCC tumour growth, likely via suppression of inflammatory/immune cell signalling and extracellular matrix deposition, but does not promote tumour angiogenesis or growth of all solid tumours.

## Introduction

Glucocorticoids are vital modulators of the physiological stress response, exerting myriad effects across a range of tissues [1]. Their potent anti-inflammatory and immunosuppressive effects have also been exploited clinically for more than half a century; synthetic glucocorticoids are commonly used to treat chronic inflammatory conditions such as rheumatoid arthritis, to suppress the immune system before organ transplant, and in the treatment of leukemia [2].

The adverse consequences of chronic glucocorticoid excess are exemplified in people with Cushing’s syndrome, who develop increased central adiposity, dyslipidemia, muscle wasting, loss of memory, hyperglycaemia and insulin resistance [1]. Reducing glucocorticoid action in key target tissues, such as liver, adipose and brain, may therefore be clinically desirable, but targeting the hypothalamic-pituitary-adrenal (HPA) axis risks compromising the systemic coordination of the stress response.

Glucocorticoids are subject to tissue-specific pre-receptor regulation by the 11β-hydroxysteroid (11β-HSD) isozymes; 11β-HSD2 converts cortisol or corticosterone to inert 11-keto metabolites (cortisone or 11-dehydrocorticosterone, respectively) to allow selective access of aldosterone to mineralocorticoid receptors (MR), while 11β-HSD1 re-activates glucocorticoids by catalyzing the reverse reductase reaction in target tissues [3], including liver, adipose, brain and the blood vessel wall [4]. Targeting 11β-HSD1 therefore offers a novel therapeutic avenue to reduce glucocorticoid action. Clinical trials of 11β-HSD1 inhibitors have shown moderate improvements in glycaemic control in patients with type II diabetes [5], and more recently have shown promise in the treatment of cognitive decline [6].

Glucocorticoids also exert potent angiostatic effects, an activity first shown over 30 years ago but the mechanism of which remains uncertain [7]. Inhibition or deletion of 11β-HSD1 promotes angiogenesis *in vitro* and *in vivo*, enhancing wound healing, reducing intra-adipose hypoxia and, most strikingly, enhancing recovery after myocardial infarction in mice [8-12]. Whilst presenting a potential clinical opportunity, these findings have also raised concerns that 11β-HSD1 inhibitors could exacerbate conditions characterised by pathological angiogenesis, such as proliferative diabetic retinopathy and solid tumour growth [15]. Whereas 11β-HSD1 inhibition or deletion was recently shown not to promote angiogenesis in a model of proliferative retinopathy [13], there is evidence to suggest it could influence tumour growth [14]. Moreover, not only might 11β-HSD1 inhibitors act in vascular cells to promote tumour angiogenesis, but they might also directly influence tumour cells as well as other cells in the tumour microenvironment, including fibroblasts, and tumour-associated immune cells [38].

The only study to address this topic thus far demonstrated that overexpression of 11β-HSD1 in hepatocellular carcinoma cells reduced tumour growth and angiogenesis [14]. No study has yet examined the effects of 11β-HSD1 inhibition on tumour growth. Of note, expression of 11β-HSD1 and the glucocorticoid receptor (GR) are particularly high in squamous cell carcinoma (SCC) [16], highlighting this tumour type as potentially glucocorticoid-sensitive. The present study tested the hypothesis that 11β-HSD1 inhibition promotes the growth of subcutaneously-implanted SCC and pancreatic ductal adenocarcinoma (PDAC) tumours in mice, as a result of increased tumour angiogenesis.

## Material and Methods

### Animals

In total, 18 C57BL6/J and 18 FVB/N mice were purchased from Envigo (Blackthorn, UK) or Charles River (Elphinstone, UK), respectively. All experimental animals were female and aged 9-14 weeks and sacrificed by cervical dislocation. Groups were age-matched. All procedures were approved by the institutional ethical committee and carried out by a licensed individual and in strict accordance with the Animals (Scientific Procedures) Act 1986 and the EU Directive 2010/63 and under project licence 70/8897 or 60/4523.

### Cell culture

Studies made use of two immortalised murine cancer cell lines. SCC cells [17] were generated in-house by Dr Alan Serrels using a two-stage 7,12-Dimethylbenz[a]anthracene (DMBA)/TPA chemical carcinogenesis protocol [18]. A PDAC cell line, Panc043, was provided by the Beatson Institute in Glasgow; these cells were originally derived from tumours developed using the *LSL-KrasG12D/+;LSL-Trp53R172H/+;Pdx-1-Cre* (KPC) model [19]. Panc-043 cells were cultured in Dulbecco’s Modified Eagle Medium (DMEM) supplemented with 10% FCS. SCC cells were maintained in Glasgow Minimum Essential Medium (GMEM) supplemented with 10% FCS, 2mM L-Glutamine, 1mM sodium pyruvate, MEM non-essential amino acids (Thermo-Fisher) and MEM vitamins.

Tumour cells (SCC or Panc043) were cultured in 96-well plates (Bio-Greiner; 5000 cells per well) and treated with 25-300nM UE2316 or corticosterone. Plates were imaged and confluence determined using the Incucyte ZOOM Live-cell analysis system (over 72 hours; Essen BioScience). An alamarBlue assay (Thermo-Fisher) was also performed as per manufacturer’s instructions to provide a secondary measure of viable cell number.

### Drugs and Corticosteroids

The 11β-HSD1 inhibitor UE2316 ([4-(2-chlorophenyl-4-fluoro-1-piperidinyl][5-(1H-pyrazol-4-yl)-3-thienyl]-methanone) was synthesised by High Force Ltd (Durham, UK) [20]. For *in vivo* studies, UE2316 was delivered *ad libitum* to animals added to a RM1 diet (175mg/kg UE2316) prepared by Special Diet Services (Essex, UK). 11-Dehydrocorticosterone and corticosterone were from Steraloids (Newport, USA). Tritiated steroids ([1,2,6,7]-^3^H_4_-corticosterone and [1,2,6,7]-^3^H_4_- cortisone) were from PerkinElmer (Wokingham, UK).

### Tumour Model

*In vivo* studies used an established model of subcutaneous tumour development [21]. SCC or PDAC cells were injected subcutaneously (1×10^6^ cells/flank) into FVB/N or C57BL6/J mice, respectively, fed either control or UE2316 diet for 5 days in advance of injection and throughout the remainder of the experiment (N=6-9/group). Diet was weighed regularly to monitor consumption, which did not differ between diets. SCC tumours were grown for 11 days, PDAC tumours were grown for 14 days, and both were measured using calipers every 2-3 days. Tumour volume was calculated as the volume of an ellipsoid (0.5*length*breadth^2^). Mice were culled by cervical dislocation.

### Histology

#### Vessel staining

Paraffin-embedded tumour sections underwent rehydration and heat-based antigen retrieval. Sections were permeabilised (0.4% Triton-X, 15 min) and blocked (1% normal goat serum, 30 min; Biosera, Nuaille, France), incubated with primary CD31 antibody (1/300 dilution, 18h, 4°C, Ab28364; Abcam, Cambridge, UK), rinsed with PBS and incubated with secondary antibody and primary conjugated α-smooth muscle actin antibody (1/1000 dilution, 1 hour, room temperature, A-11034; Molecular Probes, Eugene, USA. C6198; Sigma) before counterstaining with DAPI (5 min) and mounting using Fluoromount G (SouthernBiotech, Cambridge, UK). Slides were imaged with an Axioscan.Z1 (Zeiss) digital slide scanner. Higher magnification images were obtained using a LSM710 confocal microscope (Zeiss). Vessels were manually counted by a blinded observer across 10 randomly selected 0.1mm^2^ fields of view, from two tumour sections spaced 50 µm apart. CD31-positive/α-SMA-negative vessels and CD31/α-SMA-positive vessels were both quantified to allow the ratio of vessels with smooth muscle coverage to be calculated. As a secondary measure of vessel density, sections stained for CD31 were quantified by Chalkley count, as described [22]. One section (three hotspots) was quantified per tumour.

#### *In vivo* tumour cell proliferation

Tumour sections were stained with Ki67 antibody (proliferation marker, 1/100 dilution, Ab155580; Abcam) as above. 2 sections/tumour were scanned at 200x magnification, the most proliferative region selected by eye, and this region then imaged at 400x magnification. Ki67-positive cells were then quantified manually per hotspot.

#### Immune/inflammatory cell staining

F4/80 (1/300, 14-4801; eBiosciences) and CD3 (1/100, Sc-20047; Santa-Cruz) staining were performed using the Leica BOND-III automated staining system and the Leica refine detection kit as per manufacturer’s instructions (Leica). Trypsin-based antigen retrieval was used for F4/80 staining, and heat-based antigen retrieval for CD3 staining. Dehydrated sections were mounted with DPX and imaged using the slide scanner. Images were segmented and stain percentage area was quantified automatically using ImageJ software.

### Enzyme activity assays

A BioRad protein DC assay (BioRad, Hemel-Hempsted, UK) was performed as per manufacturer’s instructions.

#### Dehydrogenase activity assay

Homogenized tumour samples were diluted in assay buffer (63g glycerol, 8.77g NaCl, 186mg ethylenediaminetetraacetic acid (EDTA), 3.03g Tris, made up to 500mL with distilled H_2_O and pH adjusted to 7.7). ^3^H_4_-Corticosterone (250nM) and NADP^+^ (2mM; Cambridge Bioscience) were added before incubation in a shaking water bath (37°C). After incubation, samples were extracted with ethyl acetate (10:1), dried under nitrogen and dissolved in 65:15:25 water/acetonitrile/methanol.

#### Reductase activity assay

C57BL6/J mouse liver was excised and sectioned. Liver pieces (5-20mg, N=6/group) were cultured in 1mL DMEM-F12 medium containing 12.5nM ^3^H_4_-cortisone and 1µM cold cortisone with either 300nM UE2316 or vehicle (final DMSO concentration 0.3%). Plates were incubated for 24 hours (5% CO_2_, 37°C). Media was extracted on Sep-Pak C-18 (360mg) cartridges (Waters, Elstree, UK), dried under nitrogen, resuspended in 200µL HPLC-grade H_2_O added, and extracted with ethyl acetate (10:1) to remove phenol red contamination, dried under nitrogen and dissolved in 60:40 water/methanol.

### Second Harmonic Generation Imaging

Type I collagen was visualized in SCC and PDAC tumours (N=6/group) by Second Harmonic Generation (SHG) microscopy. A pump laser (tuned to 816.8 nm, 7 ps, 80 MHz repetition rate; 50 mW power at the objective) and a spatially overlapped second beam, termed the Stokes laser (1064 nm, 5–6 ps, 80 MHz repetition rate, 30 mW power at the objective; picoEmerald (APE) laser) was inserted into an Olympus FV1000 microscope coupled with an Olympus XLPL25XWMP N.A. 1.05 objective lens with a short-pass 690 nm dichroic mirror (Olympus). The Second Harmonic Generation signal was filtered (FF552-Di02, FF483/639-Di01 and FF420/40) and images quantified using Image J.

### qPCR

Frozen tissue was homogenized in Qiazol reagent (Qiagen), allowed to settle at room temperature for 5 min, vortexed in chloroform and left to settle for 2 min before centrifugation (12000 RCF x 15 min at 4°C). The resultant aqueous phase was mixed with an equal volume of 70% ethanol. All subsequent on-column steps were performed as per the RNeasy manufacturer’s protocol. RNA concentration and integrity were assessed using the Nanodrop 1000 (Thermo-Fisher Scientific). cDNA was generated from RNA using the QuantiTect Reverse Transcription Kit (Qiagen) as per manufacturer’s protocol. For the PCR reaction, samples were incubated at 42° for 15 min followed by 95° for 3 min in a Thermal cycler (Techne-Cole-Palmer, Staffordshire, UK). cDNA was diluted 1/40 in RNase-free water and a standard curve constructed by serial dilution of a pooled sample. In triplicate on a 384-well plate, 2μL of sample were combined with 5µL of Lightcyler 480 Probes Master mastermix (Roche), primers (0.1μL/sample Forward and Reverse), probe (0.1μl/sample) and RNase-free water to make up to 10μL total volume. Plates were spun (420 RCF x 2 min on LCM-3000 plate centrifuge (Grant Instruments, Royston, UK) before analysis on the Light Cycler 480 (Roche). Samples were run for 50 cycles (10s at 95°C and 30s at 60°C). All data were normalised to the average of two housekeeping genes (*Gapdh* and *Tbp*).

### RNA sequencing

RNA from SCC tumours, extracted as described above, was sequenced by GATC Biotech (Constance, Germany). Raw data were processed using Tophat2 [23], which was used to map reads to the mouse mm10 reference genome. Differential gene expression was analysed using Cuffdiff [24]. DEseq2 was used to perform a Principle Component Analysis (PCA) to assess variance between samples. Gene ontology analysis was performed using the Database for Annotation, Visualization and Integrated Discovery (DAVID) v6.8.

### Data analysis and statistics

All statistics were performed using Prism software v6/7 (Graphpad). Data are presented as mean ± S.E. Outliers were identified using Grubbs’ test and excluded appropriately. All data sets were tested for a parametric distribution and transformed/analysed appropriately. N refers to the number of animals per group in an experiment, with the exception of cell culture studies in which N refers to biological repeats on separate days using the same cell line. P<0.05 was considered significant.

## Results

### 11β-HSD1 is expressed in SCC but not PDAC tumour cell lines

When comparing 11β-HSD1 dehydrogenase activity between tumour types, SCC tumours showed a considerably higher rate of product formation than PDAC (Fig. 1A) and showed higher GR expression (Fig. 1B). 11β-HSD2 was not detected in either tumour type.

**Figure 1.**
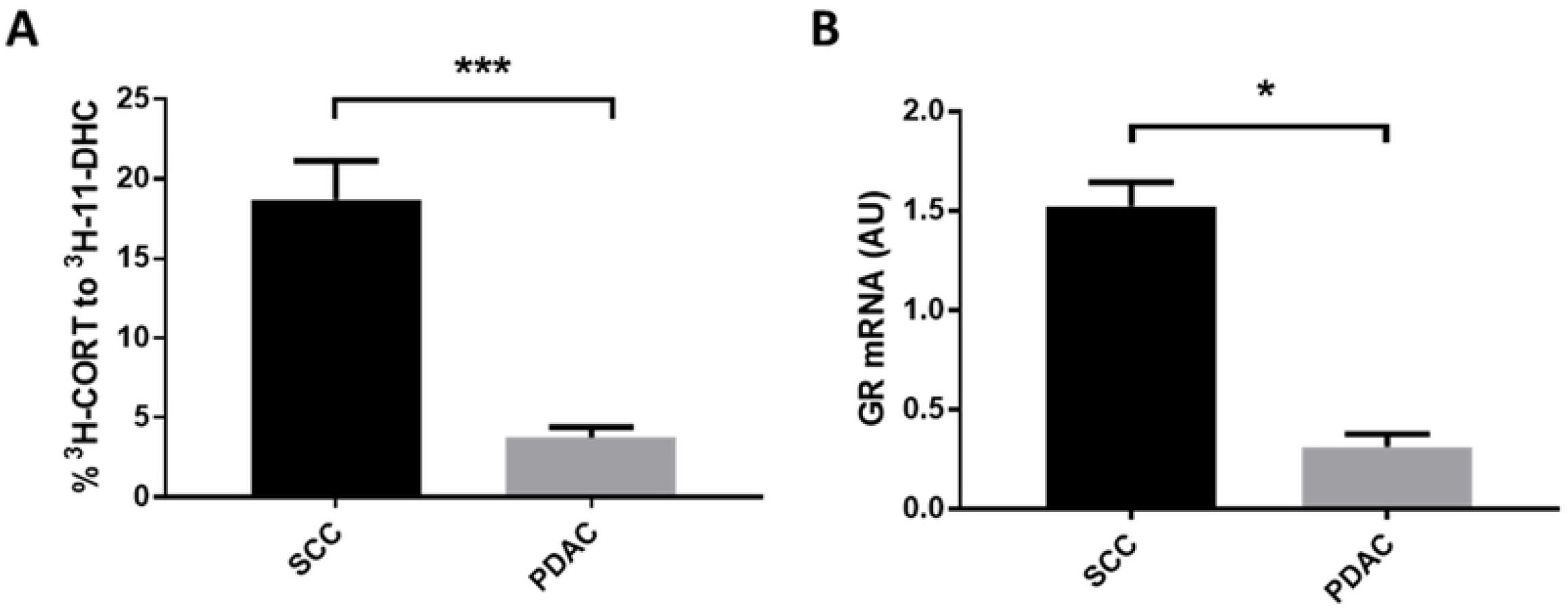
SCC tumours show greater 11β-HSD1 activity and express more GR than PDAC tumours. A) SCC tumours had greater 11β-HSD1 dehydrogenase activity than PDAC tumours. *** P<0.001. N=6/group. B) GR transcript levels were greater in SCC tumours than PDAC tumours. N=5-6/group. * P<0.05. Data were compared by independent sample t-test.

### 11β-HSD1 inhibition enhances SCC tumour growth

UE2316 accelerated the growth of SCC tumours from day 4 onwards (Fig. 2A) but had no effect on the growth of PDAC tumours (Fig. 2B). UE2316 and control diet fed groups consumed similar quantities of diet and did not differ in weight throughout the experiment (Fig. 2C). The estimated dosage achieved in the present studies (based on diet consumed per cage per 2-3 days) was 25-30mg/kg/mouse/day. Ki67 staining revealed a trend towards reduced tumour cell proliferation in UE2316-treated SCC tumours compared to control tumours, but this did not reach significance (Fig. 2D-F).

**Figure 2.**
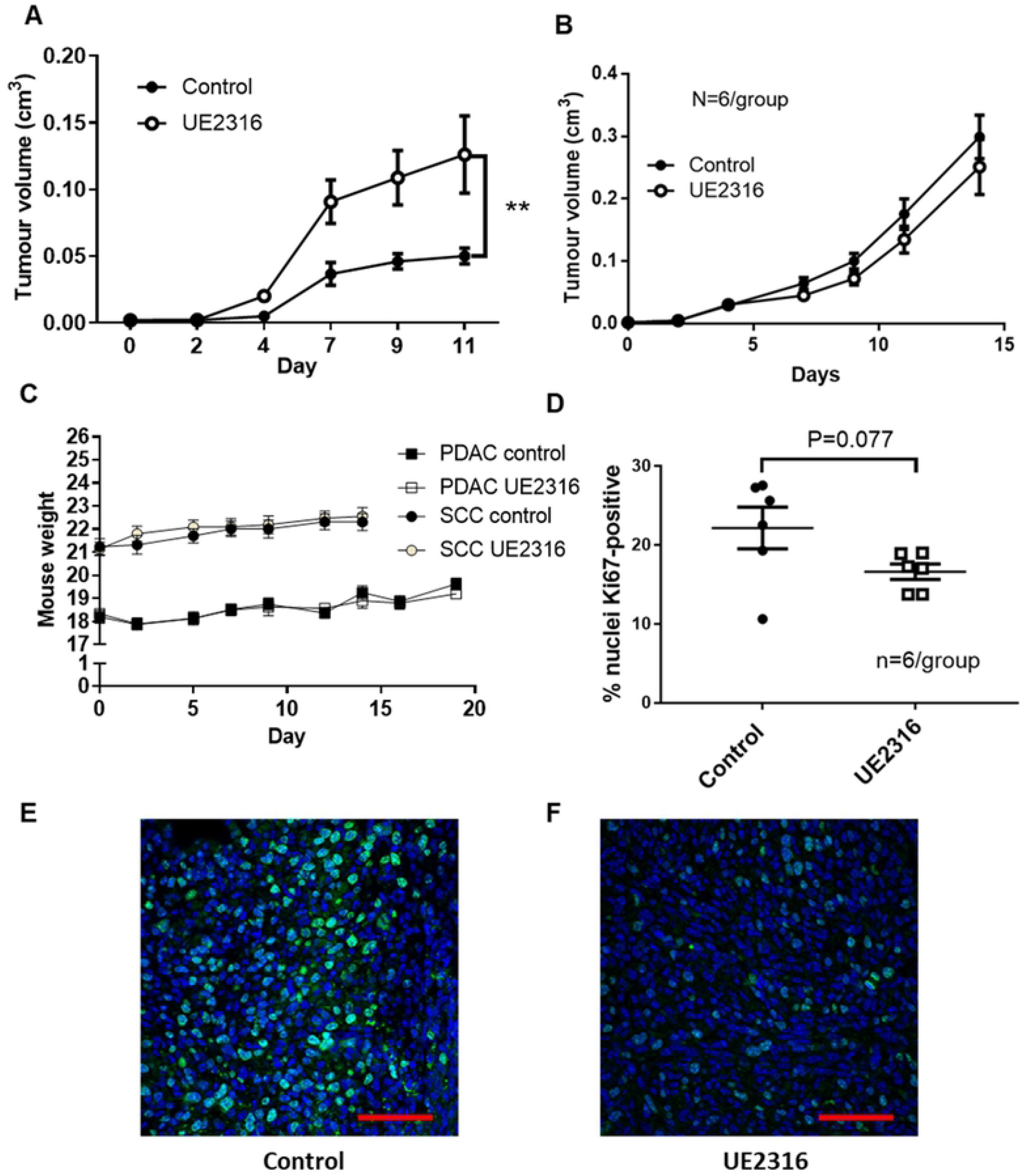
The 11β-HSD1 inhibitor UE2316 enhances SCC but not PDAC tumour growth. A) UE2316 enhanced tumour growth from day 4 onwards in mice injected with SCC cells. N=9/group. B) UE2316 did not affect PDAC tumour growth in mice injected with Panc043 cells. N=6/group. C) Neither tumour cell injection (day 5) nor UE2316 diet introduction affected mouse weight. N=6-9/group. ** P<0.01. Data were compared by 2-Way ANOVA. D) The proportion of cells staining positive for proliferation marker Ki67 showed a trend towards being reduced (P=0.07) in tumours from UE2316-treated mice but this did not reach significance. N=6/group. Data were compared by independent samples t-test. Representative images of hotspots from Ki67-stained squamous cell carcinoma (SCC) tumours from control (E) and UE2316 treated (F) mice are shown. Hotspots were typically near the periphery of the tumour. Scale bar = 50µm.

### 11β-HSD1 inhibition does not promote angiogenesis in SCC tumours

The effect of 11β-HSD1 inhibition on vessels in tumours was assessed by CD31/α-SMA-positive staining (Fig. 3A and B). In SCC and PDAC tumours, UE2316 did not affect the number of blood vessels per field of view (Fig. 3C) or the vessel number determined by Chalkley counts (Fig. 3E). In SCC tumours, UE2316 did not affect the proportion of immature vessels lacking smooth muscle coverage, assessed by CD31 staining in the absence of α-SMA staining (Fig. 3D). UE2316 also did not affect mRNA levels for angiogenic factors *Vegfa* and *Vegfr2* in either tumour type (Fig. 3F).

**Figure 3.**
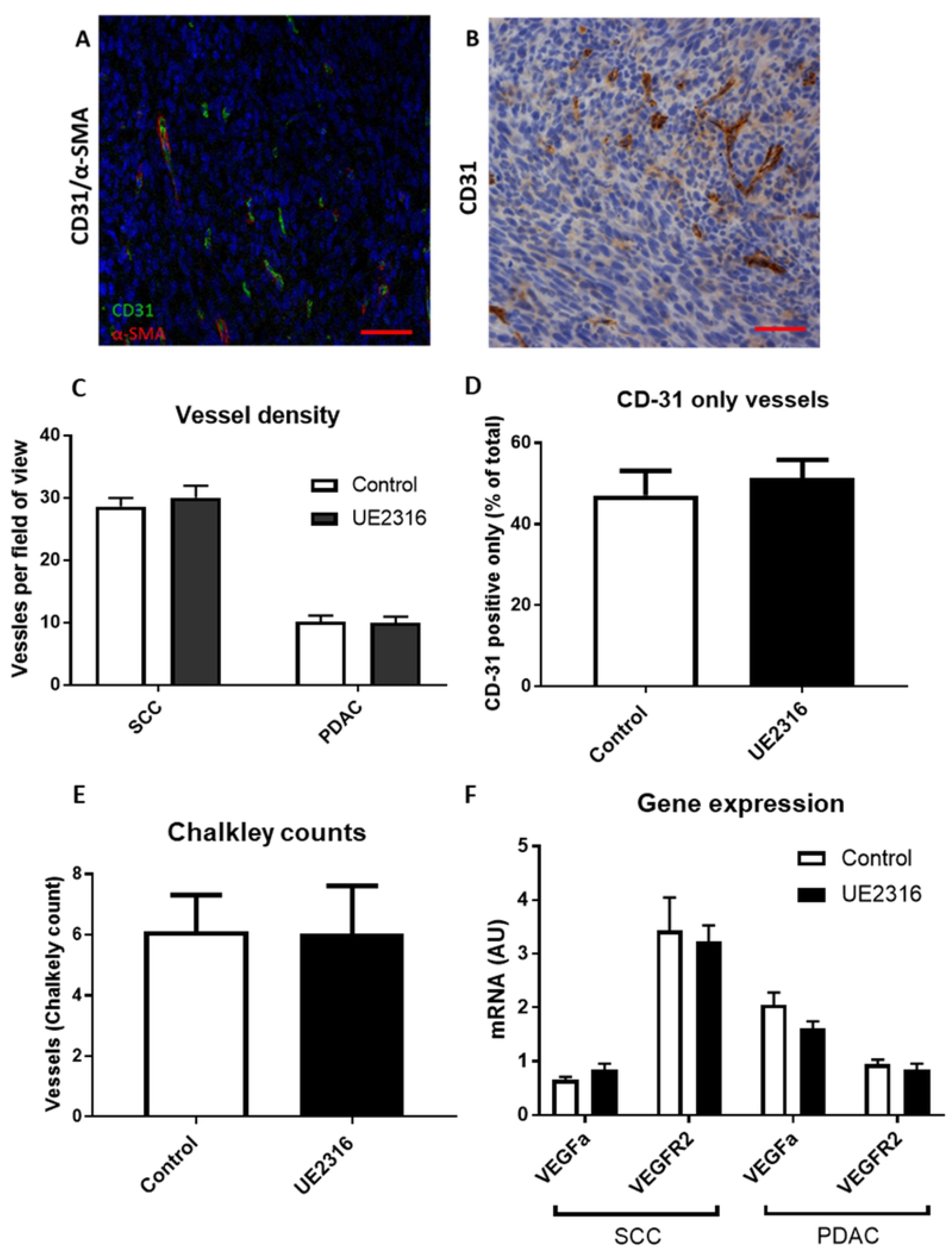
UE2316 does not affect angiogensis in tumours. A) Tumour tissue from SCC tumours; endothelial cells are stained green (CD31 visualised with Alexa-Fluor 488), smooth muscle cells are stained red (α-SMA visualised with Cy3) and nuclei are stained blue (DAPI). Tumours had densely packed nuclei. 200x magnification. Scale bar 50µm. B) CD31 was also visualised with diaminobenzidine (DAB) for counts. 200x magnification. Scale bar 50µm. C) UE2316 did not affect vessel density in either SCC or PDAC tumours. D) UE2316 did not affect the proportion of vessels lacking smooth muscle coverage in SCC tumours (i.e. CD31 positive but α-SMA negative). (E) UE2316 did not affect Chalkely counts in SCC tumours. 1 section/tumour, N=5-6 animals/group. F) mRNA levels for *Vegfa* and *Vegfr2* in SCC tumours were unaffected by UE2316. Data were compared by independent samples t-test for panels C/D, Mann-Whitney U test for Panel E.

### Neither corticosterone nor UE2316 affect SCC cell proliferation *in vitro*

SCC cells in culture were imaged using the Incucyte ZOOM live cell imaging system to investigate the effects of glucocorticoids and UE2316 on cell growth and morphology. Addition of increasing concentrations of corticosterone (Fig. 4A) or UE2316 (Fig. 4B) had no effect on the growth of SCC cells over 72 hours. Neither corticosterone (Fig. 4C) nor UE2316 (Fig. 4D) affected cell viability at any concentration, assessed after 72 hours using an alamarBlue assay.

**Figure 4.**
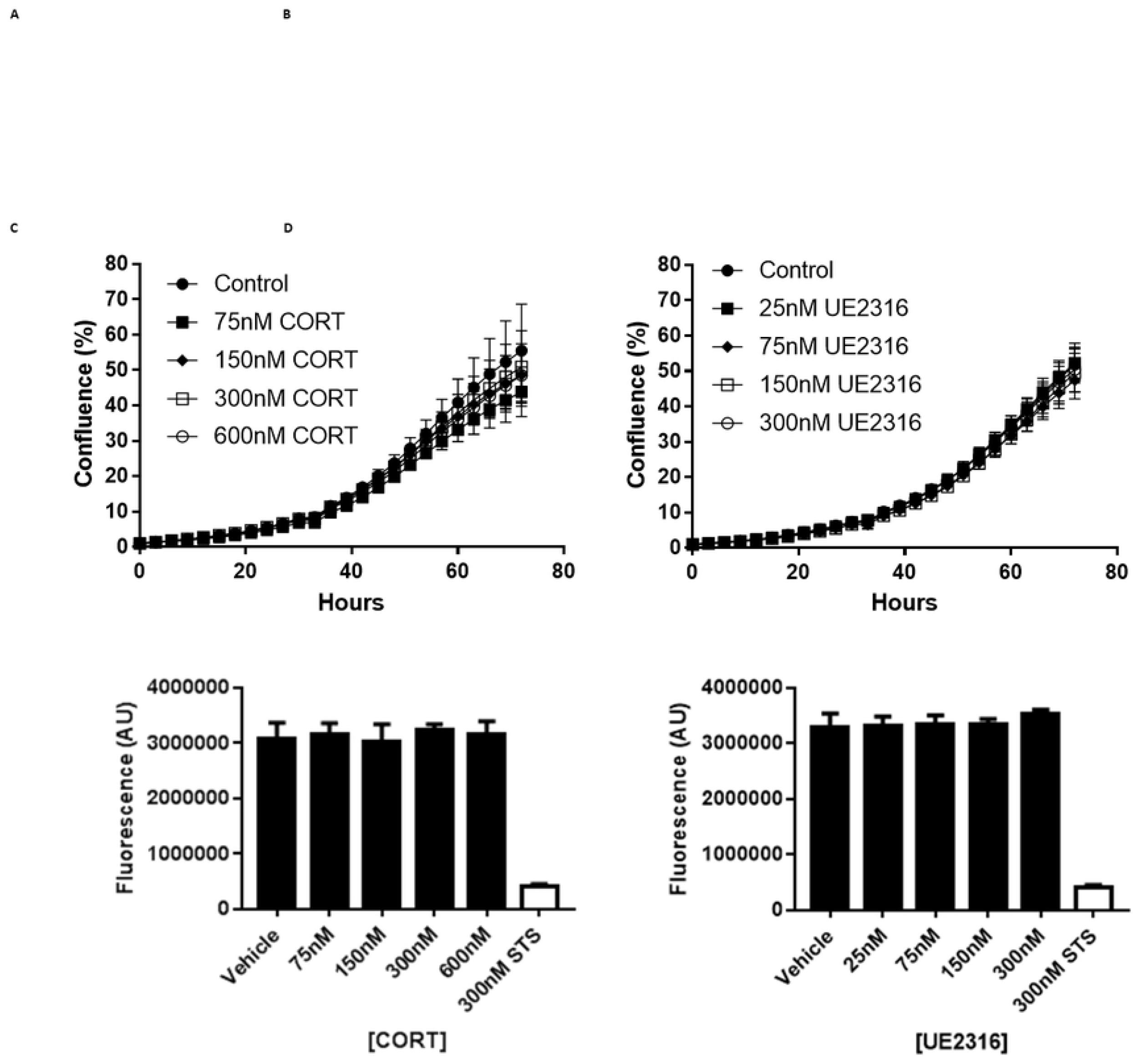
Neither corticosterone nor UE2316 affect SCC cell growth or viability *in vitro*. The confluence of SCC cells imaged over 72 hours using the Incucyte was unaffected by exposure to either corticosterone (CORT, panel A) or the 11β-HSD1 inhibitor UE2316 (panel B). 300nM STS was included in all experiments as a positive cytotoxic control. N=5 (technical repeats, treatments in sextuplet). SCC viability, as determined by the alamarBlue assay, was unaffected by the addition of corticosterone (panel C) or the 11β-HSD1 inhibitor UE2316 (panel D). AU = Arbitrary units. N=4 (technical repeats, treatments in sextuplet). Data were compared by one-way ANOVA.

### 11β-HSD1 inhibition does not alter F4/80- or CD3-positive cell infiltration into SCC tumours

To quantify inflammatory cell content, sections from SCC tumours from control (Fig. 5A) and UE2316-diet-fed mice (Fig. 5B; N=6/group) were labelled with F4/80 antibody, a macrophage marker. The antibody produced a cytoplasmic stain, present across the tumour but concentrated at the tumour periphery and in regions near the centre of the tumour. There was no significant difference in F4/80-positive area in tumours from RM-1 and UE2316 diet-fed mice, despite a trend towards a decrease in UE2316-treated tumours (Fig. 5C). To quantify infiltrating T-cells, SCC tumours from control and UE2316-diet-fed mice (N=5/group) were labelled with anti-CD3 to identify CD3-positive cells. There was no significant difference in CD3-positive area in tumours from RM-1 and UE2316 diet-fed mice (Fig. 5D); representative images are shown in Figure 5E/F.

**Figure 5.**
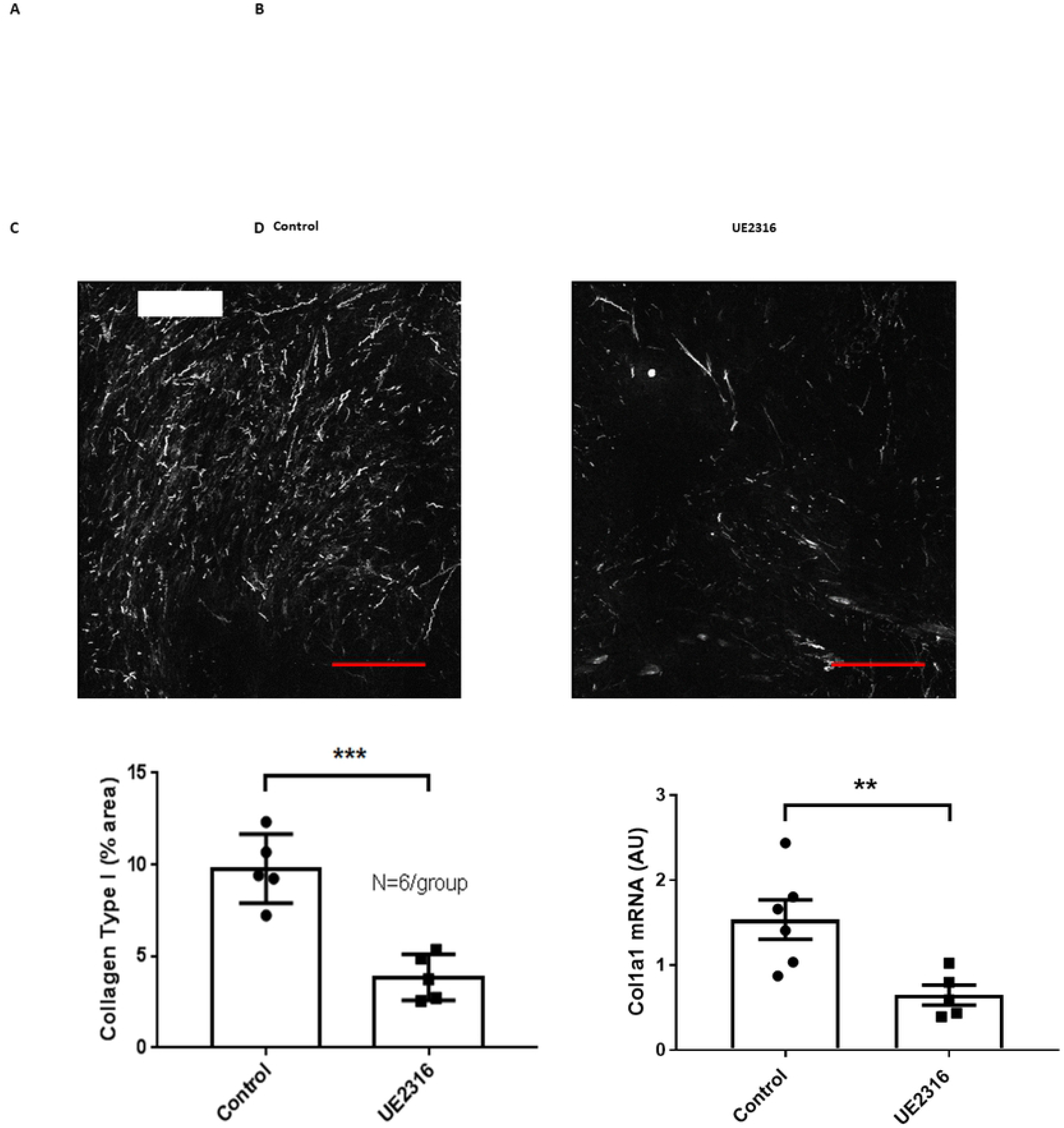
F4/80 and CD3 positive cell number in SCC tumours were unaffected by UE2316. Representative images of squamous cell carcinoma (SCC) tumours from control (A) and UE2316-treated (B) mice are shown, with DAB immunoreactivity to anti-F4/80 antibody shown in brown and haematoxylin-counterstained nuclei in blue. C) Immunostaining did not reveal a difference in F4/80-positive stain area between tumour from control and UE2316-treated mice (P=0.17). N=5- 6/group. Data were compared by independent samples t-test. Scale bar = 50µm. Immunostaining found no difference in CD3-positive stain area between SCC tumours from control and UE2316-treated mice, assessed by whole section analysis. N=5/group (D). Representative images of CD3 labelled SCC tumour sections from control (E) and UE2316-treated (F) squamous cell carcinoma (SCC). DAB immunoreactivity shown in brown and haematoxylin-counterstained nuclei in blue. Data were compared by independent samples t-test, N=5/group.

### 11β-HSD1 inhibition reduces type 1 collagen deposition in SCC tumours

To determine whether tumour collagen deposition was altered by 11β-HSD1 inhibition, Second Harmonic Generation (SHG) microscopy was performed on SCC tumours (N=6/group; Fig. 6A and B). Automatic quantification of % collagen area from SHG images revealed a reduced amount of type I collagen in tumours from mice fed UE2316-diet compared to tumours from mice fed normal diet (Fig. 6C). This difference was also apparent at a transcriptional level (Fig. 6D).

**Figure 6.**
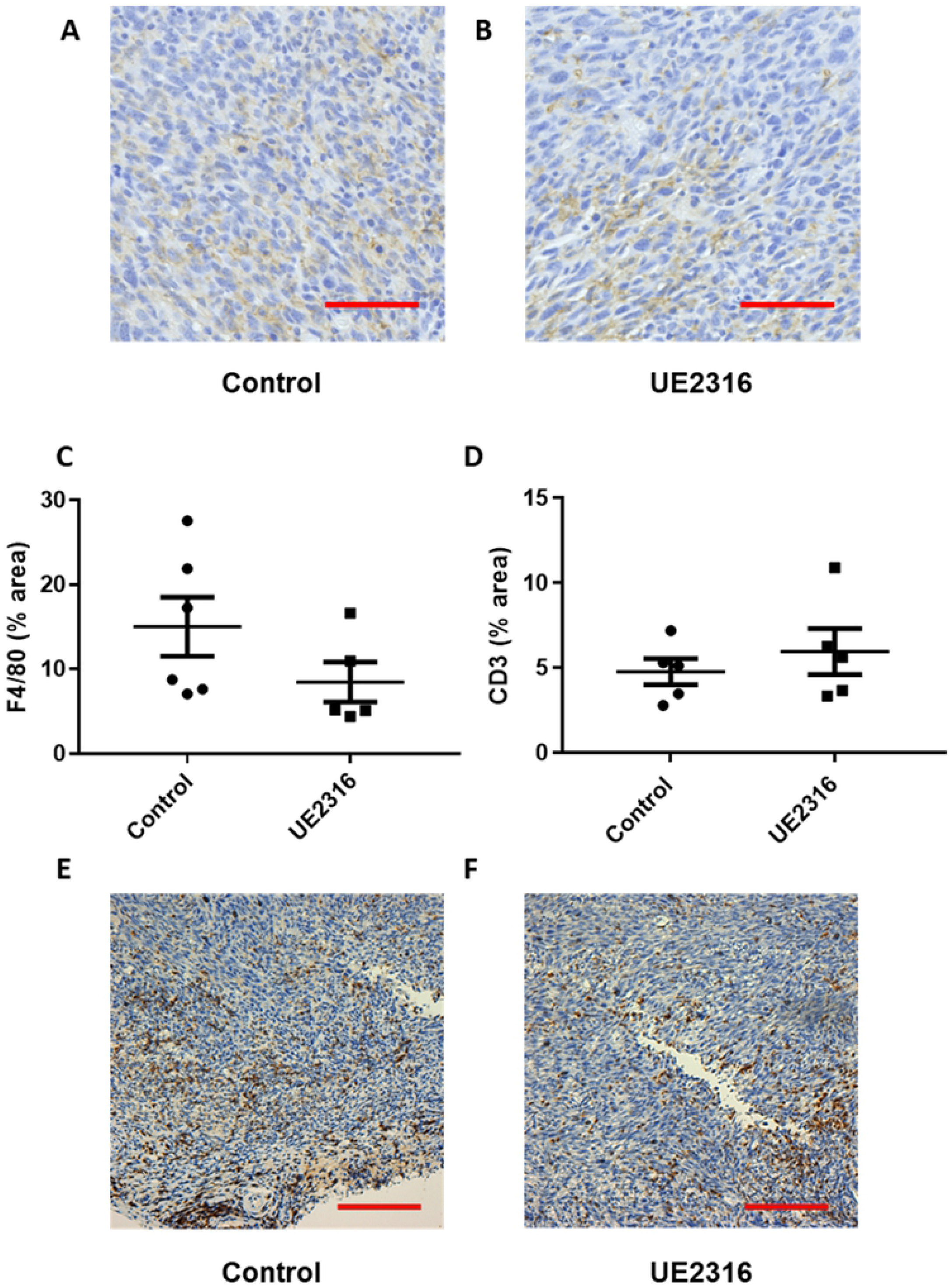
Type I collagen is reduced in SCC tumours from UE2316-treated mice. Second Harmonic Generation imaging showed type I collagen (white signal) in SCC tumours from UE2316-treated (B) and control mice (A). Scale bar = 100µm. C) Type I collagen was reduced in tumours from UE2316-treated mice. *** P<0.001. N=5/group. D) *Col1a1* mRNA was reduced in SCC tumours from UE2316-treated mice compared to control mice. AU = Arbitrary units. ** P<0.01. N=5-6/group. Data were compared by independent samples t-test.

### 11β-HSD1 inhibition influences immune and inflammatory signaling in SCC tumours

Genes found to be differentially expressed (DE) between control and UE2316-treated SCC tumours by RNA sequencing were analysed using the Database for Annotation, Visualization and Integrated Discovery (DAVID) v6.8. 674 genes were found to be differentially regulated between treatment groups. Significantly relevant (P<0.05) biological processes are listed in Table 1. Given the importance of local glucocorticoids in regulating inflammation, and the chronic inflammatory state of the tumour microenvironment, genes associated with the inflammatory response and immune response (6.1% and 4.1% of DE genes respectively, as identified by DAVID) and their relative expression in UE2316-treated tumours (as identified by RNA-seq) are shown in Figures 7A and B. mRNA coding for a large number of pro-inflammatory cytokines were reduced after UE2316 treatment, while mRNA for cytokine receptors, toll-like receptor and mast cell protease transcript was increased.

**Table 1.**

| Process | q-value |
| --- | --- |
| Inflammatory response | 3E-08 |
| Cellular response to interferon-beta | 4E-07 |
| Positive regulation of gene expression | 3E-04 |
| Immune response | 6E-04 |
| Response to lipopolysaccharide | 0.001 |
| Positive regulation of cell migration | 0.001 |
| Chemokine-mediated signalling pathway | 0.004 |
| Cellular response to interferon-γ | 0.004 |
| Negative regulation of osteoblast differentiation | 0.003 |
| Chemotaxis | 0.008 |
| Collagen fibril organization | 0.008 |
| Defence response to virus | 0.007 |
| Positive regulation of apoptotic process | 0.008 |
| Circadian rhythm | 0.009 |
| Osteoblast differentiation | 0.01 |
| Negative regulation of cell migration | 0.03 |
| Cell adhesion | 0.04 |
| Ossification | 0.04 |
| Immune system process | 0.05 |
| Positive regulation of osteoblast differentiation | 0.05 |
| Defense response to protozoan | 0.05 |

Gene ontology analysis of RNA-seq data demonstrates the importance of immune and inflammatory responses in the effects of UE2316. Gene Ontology (GO) analysis was performed using the Database for Annotation, Visualization and Integrated Discovery (DAVID) v6.8. Significance is expressed using the p-value corrected for multiple hypothesis testing using the Benjamini-Hochberg method (q-value). The listed biological processes had $q < 0.05$ .

**Figure 7.**
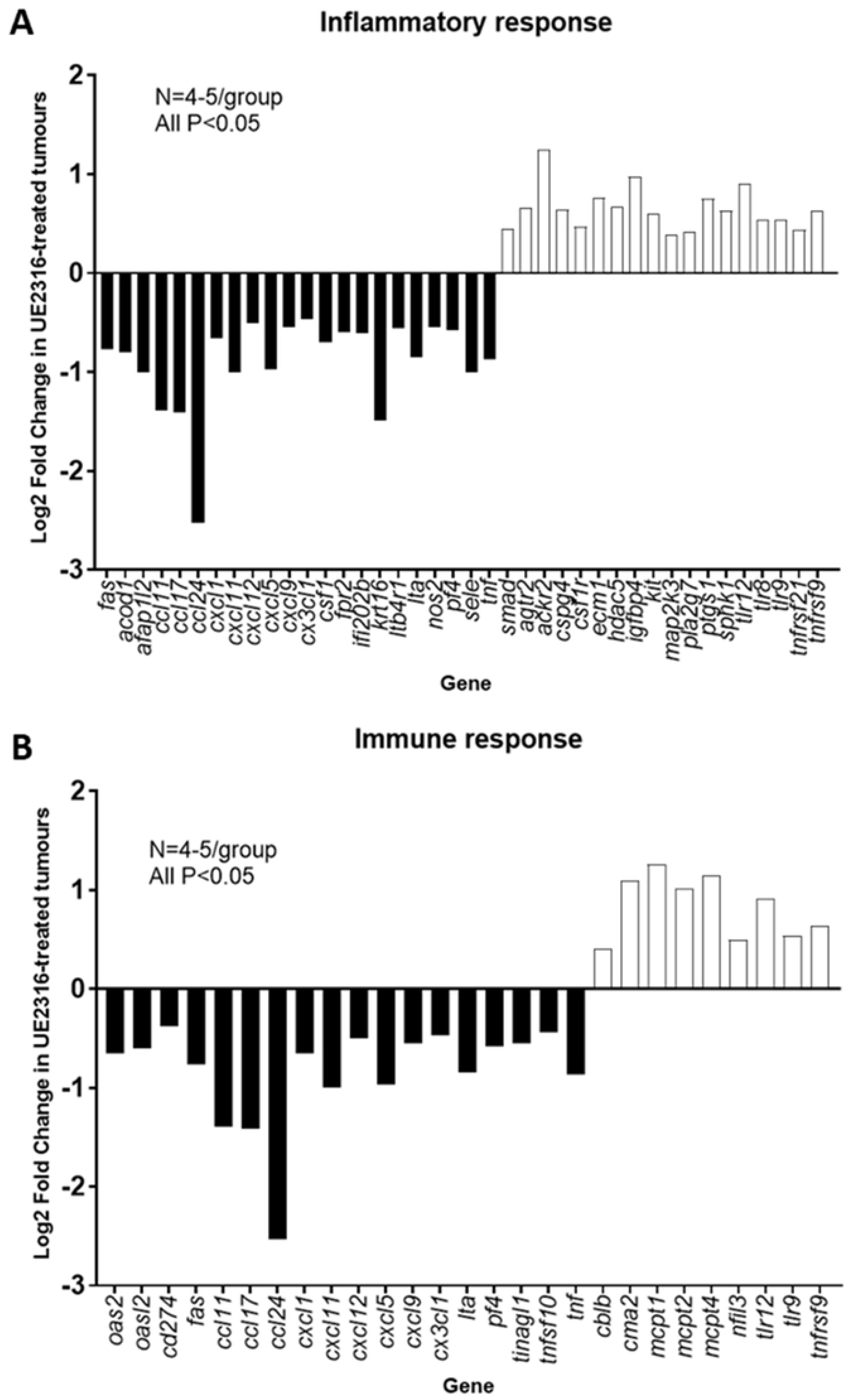
UE2316 affects inflammatory and immune response genes in SCC tumours; analysed using Gene Ontology analysis. Differentially-expressed genes identified by RNA-sequencing were defined as being related to the inflammatory response (A) or immune response (B) by Gene Ontology analysis. N=4-5/group. A modified Fisher Exact test was used to determine whether the proportion of genes in a given list was significantly associated with a biological process compared to the murine genome: P<0.05 for all the above. Data represent mean values with black bars representing genes that are down-regulated in the UE2316-treated tumours and open bars those that are up-regulated.

## Discussion

The data generated in this investigation demonstrate that 11β-HSD1 inhibition can promote SCC tumour growth in mice. This effect was not seen in PDAC tumours, which expressed lower levels of both GR and 11β-HSD1 than SCC. The present findings are in agreement with other studies that report SCC express particularly high levels of GR [25-26], suggesting that SCC may be a more glucocorticoid-sensitive tumour type than PDAC. 11β-HSD1 inhibition did not alter vessel density or *in vitro* tumour cell proliferation, but immune and inflammatory signalling pathways were altered at the transcriptomic level, as was 11β-HSD1 itself. Immune and inflammatory cell content did not differ between control and UE2316-treated SCC tumours, suggesting perhaps that cell behaviour (cytokine environment/activation state) is altered by UE2316. Fluorescence-Associated Cell Sorting of tumours would be required to more elegantly investigate this question in future studies. Generation of Type 1 collagen was reduced at both the transcriptomic and protein level in UE2316-treated SCC tumours; whether this change relates to the altered inflammatory environment remains uncertain.

The only previous study to directly manipulate 11β-HSD1 expression in a solid tumour model demonstrated that 11β-HSD1 overexpression reduced the growth of hepatocellular carcinoma (HCC) tumours in Balb/C nude mice [14], an effect which was apparent over a similar time course as the effect of 11β-HSD1 inhibition shown here (i.e. 3-5 days after cell injection). While the present study supports a role for 11β-HSD1 and local glucocorticoid metabolism in regulating tumour growth from an early stage, a different mechanism may be responsible; the study in HCC identified a significant reduction in tumour angiogenesis, attributed to reduced glycolysis, in tumours overexpressing 11β-HSD1 compared to controls. No evidence of such a process was seen in SCC tumours. The present study made use of a murine tumour cell line able to grow in mice with a functional immune system, a significant strength given that 11β-HSD1 deletion reduces T-cell infiltration in some inflammatory models [27-29] and is likely to influence the tumour microenvironment [30]. Interestingly, both tumour types used in these separate studies (SCC and HCC) were derived from tissues in which 11β-HSD1 is known to play a regulatory role (skin and liver) [31-35].

However, the present study found no evidence of enhanced angiogenesis after 11β-HSD1 inhibition. Enhanced angiogenesis and recovery post-myocardial infarction have been demonstrated consistently in 11β-HSD1 knockout mice [8, 9, 11-12) and following exposure to the 11β-HSD1 inhibitor UE2316 [36]. The reparative response to myocardial infarction is characterised by increased neutrophil and macrophage recruitment into the myocardium after 11β-HSD1 inhibition [9, 12], an effect absent in SCC tumours. Angiogenesis after induced myocardial infarction in rodents is a beneficial process and distinct from the aberrant non-resolving hypoxia-driven angiogenesis seen in tumours [37], which may be mediated by different mechanisms and explain the context-specific effects of 11β-HSD1 inhibition.

Given the lack of evidence that exaggerated angiogenesis promotes SCC tumour growth following 11β-HSD1 inhibition, we considered other mechanisms. The absence of a glucocorticoid-mediated effect on SCC cell proliferation, or any direct effect of UE2316 *in vitro*, strongly suggests that direct proliferative effects on tumour cells are not relevant. However, based on the gene ontogeny analysis of SCC tumours, the immune and inflammatory responses are likely to be of mechanistic importance. SCC tumours from UE2316-treated mice showed reduced expression of a range of pro-inflammatory cytokine and chemokine genes. These changes were accompanied by an increase in the expression of several members of the *Tlr* and *Tnfrsf* families, and *Csf1r*, suggesting reduced pro-inflammatory ligand binding. Furthermore, expression of a number of interferon-γ (IFN-γ) inducible genes was reduced in tumours from UE2316-treated animals. As TLR activation can stimulate the production of IFNs, interleukins and TNF by myeloid and lymphoid cells [38], the evidence points towards reduced inflammatory and immune cell signalling within tumours from UE2316-treated mice. The reduced expression of *Ccl* and *Cxcl* chemokines would predict reduced migration of eosinophils, neutrophils and T-cells into tumours, whilst the reduced expression of 11β-HSD1 itself after UE2316 treatment is indicative of reduced immune/inflammatory cell infiltration and activation as the enzyme is expressed in macrophages and lymphocytes and upregulated by immune cell activation [3].

The role of inflammation in tumour progression is controversial in that it can both promote tumour progression (including via stimulation of angiogenesis) and inhibit tumour progression (via anti-tumour immunosurveillance). In the present model, 11β-HSD1 inhibition appears to decrease inflammatory signalling whilst enhancing tumour growth, raising the intriguing possibility that UE2316 dampens the anti-tumour immune response. This requires confirmation at the cellular level.

11β-HSD1 inhibition has been shown to influence inflammation previously, but its effects are context-dependent and may vary between acute or chronic inflammation. Similar to induced myocardial infarction, 11β-HSD1 deficiency increases acute inflammation in models of arthritis, peritonitis and pleurisy [39-40]. In obese adipose tissue and atherosclerotic plaques from 11β-HSD1 deficient animals, however, inflammatory and immune cell infiltration is attenuated [27, 29]. Arguably, the chronic, non-resolving inflammation and hypoxia seen in obese adipose tissue and atheroma are more similar to the tumour microenvironment than to the ischaemic myocardium; thus mechanistically the latter models may be more relevant.

There is analogous evidence from other models that 11β-HSD1 influences the same inflammatory pathways as we observed here. Wamil *et al*. [27] reported that 11β-HSD1 deletion reduces similar cytokines (including members of the CCL, CXCL and TNF families) in adipose tissue from high-fat diet-fed mice, associated with decreased CD8+ T-cell infiltration and macrophage infiltration in adipose tissue. Michailidou *et al*. [10] found decreased fibrosis in adipose tissue from 11β-HSD1 knockout animals. Furthermore, 11β-HSD1 deletion reduces macrophage and T-cell infiltration into atherosclerotic plaques [29]. Several of the key gene expression changes seen in the present study have also been seen in atherosclerotic plaques after 11β-HSD1 inhibition [28], including reductions in interleukins, toll-like receptors, STAT family members, and several chemokines. The selective 11β-HSD1 inhibitor BVT-2733 was previously shown to improve symptoms of collagen-induced arthritis by reducing the expression of pro-inflammatory cytokines, including TNF, IL-1β and IL6, and reducing inflammatory cell infiltration into joints [49]. Furthermore, the beneficial effects of 11β-HSD1 inhibition in the synovium have been linked to reduced glucocorticoid action in synovial fibroblasts and osteoclasts resulting in a net reduction in damaging inflammation [50]. Although we found effects of 11β-HSD1 inhibition on transcripts in SCC tumours, these were not reflected in demonstrable differences in cell content. Staining for the T cell marker CD3 did not identify a difference between SCC tumours from control and UE2316-treated mice. Likewise, F4/80 staining did not reveal a marked difference in macrophage numbers between treatment groups, yet key transcripts for markers of macrophage polarisation were altered in whole tumour homogenates, suggesting a more subtle effect of 11β-HSD1 inhibition on macrophage content or polarisation.

Cancer-associated fibroblasts and extracellular matrix (ECM) deposition can also influence tumour progression [41-43]. The reduced type 1 collagen seen in SCC tumours mirrors the reduced fibrosis in obese adipose tissue from 11β-HSD1 deficient mice [15], which also showed decreased alpha-smooth muscle actin expression, suggesting reduced fibroblast numbers. Reduced fibrosis and reduced expression of *Col1a1, Col1a2, Col14a1*, stromal-cell derived factor 1 (*Sdf1*) and *Lox* are all suggestive of reduced fibroblast activity [41-42]. Since fibroblasts can promote anti-tumour immune cell infiltration into tumours [41, 43], suppression of fibroblasts by UE2316 could explain the potentially dampened anti-tumour immune response in SCC tumours. Conversely, inflammatory cells are also able to recruit fibroblasts into SCC tumours [44] and this enhanced recruitment can promote SCC growth suppression via the deposition of a fibrotic ECM that constrains tumour cell proliferation and invasiveness [45-46], so the effect of UE2316 could be primarily on inflammatory cells or on tumour cells releasing pro-inflammatory signals, with secondary effects on fibroblasts.

Given that only one of the two tumour types examined responded to UE2316 treatment, predicting which tumour types may be more at risk will be important if 11β-HSD1 inhibitors are to be used in at-risk patients. Review of cancer genomics data sets available via the cBioPortal for Cancer Genomics [51] reveals amplification of *HSD11B1* expression in 8-10% of breast and hepatobiliary cancer studies, while around 8% of cutaneous melanomas show either mutation (4%) or amplification (4%) of the gene. Altered expression of *HSD11B1* is also apparent in around 5% of studies on endometrial cancers, non-Hodgkin lymphomas, non-small cell lung cancers and melanomas. Extra vigilance is recommended if 11β-HSD1 use is indicated in patients with such *HSD11B1*-expressing tumours.

In summary, inhibition of 11β-HSD1 in SCC tumours does not alter tumour angiogenesis but dampens immune and inflammatory signalling within the tumour microenvironment, possibly leading to the reduced activation of cancer associated fibroblasts and the reduced deposition of type I collagen. These factors, in combination, may promote SCC growth in this model but relevance to other tumours is uncertain.

## Acknowledgments

*CTD, EM, MM, JCD, ML and SA designed and conducted experimental work; AS, SPW, NZMH, and RA provided tools and supervised specialised analyses; VGB, PWFH and BRW supervised CDT in experimental design, analysis and interpretation and in the preparation of the manuscript. All authors reviewed the final manuscript. Thea authors are grateful to Marisa Magennis for administrative support. Our work is supported by the British Heart Foundation and The Wellcome Trust*.

## Notes

### Competing Interest Statement

Brian Walker, Patrick Hadoke and Scott Webster are inventors on relevant patents owned by the University of Edinburgh and Brian Walker, Scott Webster and Ruth Andrew have received consultancy fees and honoraria from several companies developing selective 11?-HSD1 inhibitors. Callam Davidson is currently employed as an Associate Editor at PLOS Medicine

## References

1. Walker, BR. Glucocorticoids and Cardiovascular Disease. European Journal of Endocrinology. 2007;157(5):545–559.

2. Coutinho, AE and Chapman, KE. The anti-inflammatory and immunosuppressive effects of glucocorticoids, recent developments and mechanistic insights. Molecular and Cellular Endocrinology. 2011;335(1):2–13.

3. Chapman, KE, Holmes, M and Seckl, J. 11β-hydroxysteroid dehydrogenases: intracellular gate-keepers of tissue glucocorticoid action. Physiological Reviews. 2013;93(3):1139–1206.

4. Seckl, JR and Walker, BR. Minireview: 11β-Hydroxysteroid Dehydrogenase Type 1— A Tissue-Specific Amplifier of Glucocorticoid Action 1, Endocrinology. 2001;142(4):1371– 1376.

5. Anderson, A and Walker, BR. 11β-HSD1 Inhibitors for the Treatment of Type 2 Diabetes and Cardiovascular Disease. Drugs. 2013;73(13):1385–1393.

6. Webster, SP, McBride A, Binnie, M, Sooy, K, Seckl, JR, Andrew, R et al. Selection and early clinical evaluation of the brain-penetrant 11β-hydroxysteroid dehydrogenase type 1 (11β-HSD1) inhibitor UE2343 (XanamemTM). British Journal of Pharmacology. 2017;174(5):396–408.

7. Folkman, J, Langer, R, Linhardt, R, Haudenschild, C and Taylor, S. Angiogenesis inhibition and tumor regression caused by heparin or a heparin fragment in the presence of cortisone. Science. 1983;221(4612):719–725.

8. Small, GR, Hadoke, PWF, Sharif, I, Dover, AR, Armour, D, Kenyon, CJ et al. Preventing local regeneration of glucocorticoids by 11beta-hydroxysteroid dehydrogenase type 1 enhances angiogenesis. Proceedings of the National Academy of Sciences of the United States of America. 2005;102(34):12165–12170.

9. McSweeney, SJ, Hadoke, PWF, Kozak, AM, Small, GR, Khaled, H, Walker, BR et al. Improved heart function follows enhanced inflammatory cell recruitment and angiogenesis in 11β-HSD1-deficient mice post-MI. Cardiovascular Research. 2010;88(1):159–167.

10. Michailidou, Z, Turban, S, Miller, E, Zou, XT, Schrader, J, Ratcliffe, PJ et al. Increased Angiogenesis Protects against Adipose Hypoxia and Fibrosis in Metabolic Disease-resistant 11β-Hydroxysteroid Dehydrogenase Type 1 (HSD1)-deficient Mice. Journal of Biological Chemistry. 2012;287(6):4188–4197.

11. White, CI, Jansen, MA, McGregor, K, Mylonas, KJ, Richardson, RV, Thomson, A et al. Cardiomyocyte and vascular smooth muscle independent 11β-hydroxysteroid dehydrogenase 1 amplifies infarct expansion, hypertrophy and the development of heart failure following myocardial infarction in male mice. Endocrinology. 2016;157(1):346–357.

12. Mylonas, KJ, Turner, NA, Bageghni, SA, Kenyon, CJ, White, CI, McGregor et al. 11β- HSD1 suppresses cardiac fibroblast CXCL2, CXCL5 and neutrophil recruitment to the heart post MI. The Journal of Endocrinology. 2017;233(3):315–327.

13. Davidson, CT, Dover, AR, McVicar, CM, Megaw, R, Glenn, JV, Hadoke, PWF et al. Inhibition or deletion of 11β-HSD1 does not increase angiogenesis in ischemic retinopathy. Diabetes and Metabolism. 2017;43(5):480–483.

14. Liu, X, Tan, X, Xia, M, Wu, C, Song, J, Wu, J et al. Loss of 11βHSD1 enhances glycolysis, facilitates intrahepatic metastasis, and indicates poor prognosis in hepatocellular carcinoma. Oncotarget. 2016;7(2):2038–53.

15. Verdegem, D, Moens, S, Stapor, P and Carmeliet, P. Endothelial cell metabolism: parallels and divergences with cancer cell metabolism.Cancer & Metabolism. 2014;2;19.

16. Azher, S, Azami, O, Amato, C, McCullough, M, Celentano, A and Cirillo, N. The Non-Conventional Effects of Glucocorticoids in Cancer. Journal of Cellular Physiology. 2016;231(11):2368–73.

17. Serrels, A, McLeod, K, Canel, M, Kinnaird, A, Graham, K, Frame, MC et al. The role of focal adhesion kinase catalytic activity on the proliferation and migration of squamous cell carcinoma cells’. International Journal of Cancer. 2012;131(2):287–297.

18. McLean GW, Komiyama NH, Serrels B, Asano H, Reynolds L, Conti F et al. Specific deletion of focal adhesion kinase suppresses tumor formation and blocks malignant progression. Genes Dev. 2004;18:2998–3003.

19. Hingorani, SR, Wang, L, Multani, AS, Combs, C, Deramaudt, TB, Hruban, RH et al. Trp53R172H and KrasG12D cooperate to promote chromosomal instability and widely metastatic pancreatic ductal adenocarcinoma in mice. Cancer Cell. 2005;7(5):469–83.

20. Webster SP, Seckl JR, Walker BR, Ward P, Pallin TD, Dyke HJ et al. 4-phenyl-piperidin-1-yl)-[5-(1H-pyrazol-4-yl)-thiophen-3-yl]-methanone Compounds and Their Use. PCT Intl WO2011/033255. 2011.

21. Serrels, A, Lund, T, Serrels, B, Byron, A, McPherson, RC, von Kriegsheim, A, et al. Nuclear FAK controls chemokine transcription, Tregs, and evasion of anti-tumor immunity. Cell. 2015;163(1):160–73.

22. Hansen, S, Grabau, DA, Sørensen, FB, Bak, M, Vach, W and Rose, C. The prognostic value of angiogenesis by Chalkley counting in a confirmatory study design on 836 breast cancer patients. Clinical Cancer Research. 2000;6(1):139–146.

23. Trapnell, C, Pachter, L, Salzberg, SL. TopHat: discovering splice junctions with RNA-Seq. Bioinformatics. 2009;25(9):1105–1111

24. Trapnell, C, Roberts, A, Goff, L, Pertea, G, Kim, D, Kelley, DR et al. Differential gene and transcript expression analysis of RNA-seq experiments with TopHat and Cufflinks. Nat Protocols. 2012;7(3):562–78.

25. Budunova, IV, Carbajal, S, Kang, H, Viaje, A, Slaga, TJ. Altered glucocorticoid receptor expression and function during mouse skin carcinogenesis, Molecular Carcinogenesis. 1997;18(3):177–85.

26. Spiegelman, VS, Budunova, IV, Carbajal, S, Slaga, TJ. Resistance of transformed mouse keratinocytes to growth inhibition by glucocorticoids. Molecular Carcinogenesis. 1997;20(1):99–107.

27. Wamil, M, Battle, JH, Turban, S, Kipari, T, Seguret, D, de Sousa Peixoto, R, et al. Novel Fat Depot-Specific Mechanisms Underlie Resistance to Visceral Obesity and Inflammation in 11 -Hydroxysteroid Dehydrogenase Type 1-Deficient Mice. Diabetes. 2011;60(4):1158–1167.

28. Luo, MJ, Thieringer, R, Springer, MS, Wright, SD, Hermanowski-Vosatka, A, Plump, A et al. 11β-HSD1 inhibition reduces atherosclerosis in mice by altering proinflammatory gene expression in the vasculature. Physiological Genomics. 2012;45:47–57.

29. Kipari, T, Hadoke, PWF, Iqbal, J, Man, TY, Miller, E, Coutinho, AE et al. 11 beta-hydroxysteroid dehydrogenase type 1 deficiency in bone marrow-derived cells reduces atherosclerosis. The FASEB Journal. 2013;27(4):1519–1531.

30. Kim, R, Emi, M, Tanabe, K. Cancer immunoediting from immune surveillance to immune escape, Immunology. 2007;121(1):1–14.

31. Tiganescu, A, Tahrani, AA, Morgan, SA, Otranto, M, Desmoulière, A, Abrahams, L et al. 11β-Hydroxysteroid dehydrogenase blockade prevents age-induced skin structure and function defects. Journal of Clinical Investigation. 2013;123(7):3051–3060.

32. Terao, M, Murota, H, Kimura, A, Kato, A, Ishikawa, A, Igawa, K et al. 11β-Hydroxysteroid Dehydrogenase-1 Is a Novel Regulator of Skin Homeostasis and a Candidate Target for Promoting Tissue Repair. PLoS ONE. 2011;6(9):e25039.

33. Terao, M, Tani, M, Itoi, S, Yoshimura, T, Hamasaki, T, Murota, H et al. 11β-hydroxysteroid dehydrogenase 1 specific inhibitor increased dermal collagen content and promotes fibroblast proliferation. PloS ONE. 2014;9(3):e93051.

34. Itoi, S, Terao, M, Murota, H, Katayama, I. 11β-Hydroxysteroid dehydrogenase 1 contributes to the pro-inflammatory response of keratinocytes. Biochemical and Biophysical Research Communications. 2013;440(2);265–270.

35. Kuo, T, McQueen, A, Chen TC, Wang, JC. Regulation of Glucose Homeostasis by Glucocorticoids. In: Wang JC, Harris C, eds. Glucocorticoid Signaling. Advances in Experimental Medicine and Biology, vol 872. New York: Springer;2015:99–126.

36. McGregor, K, Mylonas, KJ, White, C, Walker, BR and Gray, G. 216 Immediate Pharmacological Inhibition of Local Glucocorticoid Generation increases Angiogenesis and Improves Cardiac Funcion after Myocardial Infarction. Heart. 2014;100 Suppl: A118.

37. Chung, AS, Ferrara, N. Developmental and Pathological Angiogenesis. Annual Review of Cell and Developmental Biology. 2011;27(1);563–584.

38. Chapman, KE, Coutinho, AE, Zhang, Z, Kipari, T, Savill, JS and Seckl, JR. Changing glucocorticoid action: 11β-hydroxysteroid dehydrogenase type 1 in acute and chronic inflammation. The Journal of Steroid Biochemistry and Molecular Biology. 2013;137(100):82–92.

39. Coutinho, AE, Gray, M, Brownstein, DG, Salter, DM, Sawatzky, DA, Clay, S et al. 11β-Hydroxysteroid dehydrogenase type 1, but not type 2, deficiency worsens acute inflammation and experimental arthritis in mice. Endocrinology. 2012;153(1);234–40.

40. Coutinho, AE, Kipari, TMJ, Zhang, Z, Esteves, CL, Lucas, CD, Gilmour, JS et al. 11β-Hydroxysteroid Dehydrogenase type 1 is expressed in neutrophils and restrains an inflammatory response in male mice. Endocrinology. 2016;157(7):2928–36.

41. Harper, J, Sainson, RCA. Regulation of the anti-tumour immune response by cancer-associated fibroblasts. Seminars in Cancer Biology. 2014;25:69–77.

42. Fang, MM, Yuan, J, Peng, C, Li, Y. Collagen as a double-edged sword in tumor progression. Tumor Biology. 2014;35:2871–2882.

43. Özdemir, BC, Pentcheva-Hoang, T, Carstens, JL, Zheng, X, Wu, CC, Simpson, TR et al. Depletion of Carcinoma-Associated Fibroblasts and Fibrosis Induces Immunosuppression and Accelerates Pancreas Cancer with Reduced Survival. Cancer Cell. 2014;25(6);719– 734.

44. Coussens, LM, Raymond, WW, Bergers, G, Laig-Webster, M, Behrendtsen, O, Werb, Z et al. Inflammatory mast cells up-regulate angiogenesis during squamous epithelial carcinogenesis. Genes & Development. 1999;13(11):1382–97.

45. Willhauck, MJ, Mirancea, N, Vosseler, S, Pavesio, A, Boukamp, P, Mueller, MM et al. Reversion of tumor phenotype in surface transplants of skin SCC cells by scaffold-induced stroma modulation. Carcinogenesis. 2006;28(3):595–610.

46. Cretu, A, Brooks, PC. Impact of the non-cellular tumor microenvironment on metastasis: Potential therapeutic and imaging opportunities. Journal of Cellular Physiology. 2007;213(2):391–402.

47. Baba, Y, Iyama, KI, Ikeda, K, Ishikawa, S, Hayashi, N, Miyanari, N et al. The Expression of Type IV Collagen α6 Chain Is Related to the Prognosis in Patients with Esophageal Squamous Cell Carcinoma. Annals of Surgical Oncology. 2008;15(2):555–565.

48. Erez, N, Truitt, M, Olson, P, Hanahan, D, Hanahan, D. Cancer-Associated Fibroblasts Are Activated in Incipient Neoplasia to Orchestrate Tumor-Promoting Inflammation in an NF-κB-Dependent Manner. Cancer Cell. 2010;17(2);135–147.

49. Zhang, L, Dong, Y, Zou, F, Wu, M, Fan, C & Ding, Y. 11β-Hydroxysteroid dehydrogenase 1 inhibition attenuates collagen-induced arthritis. International Immunopharmacology. 2013;17(3):489–94.

50. Hardy, RS, Seibel, MJ, Cooper, MS. Targeting 11β-hydroxysteroid dehydrogenases: a novel approach to manipulating local glucocorticoid levels with implications for rheumatic disease, Current Opinion in Pharmacology. 2013;13(3):440–4.

51. Cerami, E, Gao, J, Dogrusoz, U, Gross, BE, Sumer, SO, Aksoy, BA et al. The cBio cancer genomics portal: an open platform for exploring multidimensional cancer genomics data. Cancer Discovery, 2012;2(5):401–404.

52. Mitić, T, Shave, S, Semjonous, N, McNae, I, Cobice, DF, Lavery, GG et al. 11β-Hydroxysteroid dehydrogenase type 1 contributes to the balance between 7-keto- and 7-hydroxy-oxysterols in vivo. Biochemical Pharmacology, 2013;86(1):146–153.

